# Glucuronoxylomannan intranasal challenge prior to *Cryptococcus neoformans* pulmonary infection enhances cerebral cryptococcosis in rodents

**DOI:** 10.1101/2022.10.24.513461

**Authors:** Hiu Ham Lee, Dylan J. Carmichael, Victoria Ríbeiro, Dana N. Parisi, Melissa E. Munzen, Mohamed F. Hamed, Ettiman Kaur, Ayush Mishra, Jiya Patel, Rikki B. Rooklin, Amina Sher, Maria A. Carrillo-Sepulveda, Eliseo A. Eugenin, Michael R. Dores, Luis R. Martinez

## Abstract

The encapsulated fungus *Cryptococcus neoformans* is the most common cause of fungal meningitis, with the highest rate of disease in patients with AIDS or immunosuppression. This microbe enters the human body via inhalation of infectious particles. *C. neoformans* capsular polysaccharide, in which the major component is glucuronoxylomannan (GXM), extensively accumulates in tissues and compromises host immune responses. *C. neoformans* travels from the lungs to the bloodstream and crosses to the brain via transcytosis, paracytosis, or inside of phagocytes using a “Trojan horse” mechanism. The fungus causes life-threatening meningoencephalitis with high mortality rates. Hence, we investigated the impact of intranasal exogenous GXM administration on *C. neoformans* infection in C57BL/6 mice. GXM enhances cryptococcal pulmonary infection and facilitates fungal systemic dissemination and brain invasion. Pre-challenge of GXM results in detection of the polysaccharide in lungs, serum, and surprisingly brain, the latter likely reached through the nasal cavity. GXM significantly alters endothelial cell tight junction protein expression *in vivo*, suggesting significant implications for the *C. neoformans* mechanisms of brain invasion. Using a microtiter transwell system, we showed that GXM disrupts the trans-endothelial electrical resistance, weakening the human brain endothelial cell monolayers co-cultured with pericytes, supportive cells of blood vessels/capillaries found in the blood-brain barrier (BBB), and promotes *C. neoformans* BBB penetration. Our findings should be considered in the development of therapeutics to combat the devastating complications of cryptococcosis that results in an estimated ∼200,000 deaths worldwide each year.

**AUTHOR SUMMARY:** *Cryptococcus neoformans* infection of the central nervous system (CNS) typically begins by inhalation of fungal spores and results in devastating mortality rates worldwide. Over 200,000 deaths have been reported annually, with cryptococcal meningoencephalitis being the most severe form of the disease. This study investigates the ability of the fungus to invade, colonize, and cause damage to the host through properties of the fungal polysaccharide capsule, which allows the microbe a variety of both protective and offensive abilities. This capsule, made primarily of the polysaccharide glucuronoxylomannan (GXM), has been implicated in the progression and severity of cryptococcal infection in the CNS. We determined that GXM increases the fungal burden in the lungs of mice and enhances fungal migration to the brain. Interaction of GXM with the blood-brain barrier, which is a protective structure that regulates movement of particles into the CNS, demonstrated that GXM can disrupt the integrity of this barrier, compromising the delicate balances of fluids, immune cells, and other factors vital to the maintenance of the CNS. The findings of this study reveal the substantial role of GXM in establishing *C. neoformans* infection in the brain and necessitate future studies to further understand these interactions.

## INTRODUCTION

*Cryptococcus neoformans* is an encapsulated opportunistic yeast-like fungus responsible for life-threatening meningoencephalitis in both immunocompromised and previously healthy individuals. Cryptococcosis is responsible for an estimated 200,000 deaths per year globally [1], especially in HIV infected patients. *C. neoformans* initially infects humans during early childhood [2] through inhalation of small fungal particles in the form of desiccated or poorly encapsulated yeast or basidiospores [3, 4]. In most immunocompetent hosts, the fungus is controlled by local host responses and remains dormant in the lungs. However, reactivation can occur by immunosuppression, resulting in fungal replication in the lungs followed by dissemination via the bloodstream or lymphatic system to other organs, especially the brain. During cryptococcosis, fungemia is detected in about 50% of HIV-infected patients [5]. The correlation between fungemia and dissemination, including brain invasion, has been utilized for the development of experimental models of cryptococcosis [6], and fungemia is identified as an independent parameter of early mycological failure in humans [5]. *C. neoformans* crosses the blood-brain barrier (BBB) [7] and bypasses brain endothelial cells either by transcytosis, paracytosis, or via a “Trojan horse” mechanism by being transported within host inflammatory cells [8]. Once the fungus reaches the brain, it can cause lethal meningoencephalitis, which is the most serious pathological manifestation of cryptococcosis.

Microbial pathogenesis depends on the ability of the host to combat foreign infection. Microbes like *C. neoformans* have specific virulence factors, including melanin production, the ability to grow at mammalian temperatures, and the polysaccharide capsule, that allow them to evade multiple immune defenses and damage the host. The capsule of *C. neoformans* is the most distinctive physical structure of the cryptococcal cell, located outside the cell wall [9, 10]. This structure may protect the fungus from changes in the environment such as desiccation or against predation by soil amoeba [11]. The capsule is composed of at least three components: glucuronoxylomannan (GXM), galactoxylomannan, and mannoprotein, with GXM being its major constituent. Genetic and molecular biology strategies have shown that the capsule is a major contributor to *C. neoformans* infection since non-encapsulated strains have reduced virulence [12]. Immunocompromised individuals with cryptococcosis typically have high levels of *C. neoformans* capsular polysaccharide (CPS) in their serum and cerebrospinal fluid [13, 14]. Furthermore, the extensive accumulation of GXM in tissue is believed to be a major contributor to *C. neoformans* pathogenesis. This compound has been associated with a variety of immunosuppressive effects [15], such as interference with phagocytosis, antigen presentation, leukocyte migration and proliferation, as well as impediment of specific antibody (Ab) responses. GXM also enhances HIV replication [16]. However, there is lack of information on the role of *C. neoformans* GXM on lung dissemination into the bloodstream and brain invasion or pathogenesis.

Immunohistochemical analysis of post-mortem individuals’ brains with cryptococcosis exhibited substantial levels of GXM released around large penetrating vessels [17] suggesting an important role of this carbohydrate on central nervous system (CNS) invasion by this fungus. In fact, *C. neoformans* sheds large amounts (range µg to mg/mL) of polysaccharide into the cerebrospinal fluid (CSF) and infected tissues [14]. In this study, we challenged C57BL/6 mice with exogenous purified GXM before pulmonary infection with cryptococci and compared these mice to animals that were not sensitized with the polysaccharide. We found that exogenous GXM reaches the CNS, exacerbates fungal pulmonary disease, enhances *C. neoformans* dissemination from the lungs to the CNS, and has detrimental effects on tight junction proteins (TJs), the molecules that regulate endothelial cells in the BBB. These observations suggest that GXM intranasal (i.n.) challenge prior to *C. neoformans* pulmonary infection enhances brain invasion and colonization in rodents via disruption of the TJs on endothelial cells of the BBB.

To investigate the disruption of BBB integrity by GXM, we measured the activation of RhoA, a member of the Ras-related small GTPase Rho family involved in the signaling pathway for cytoskeletal regulation [18]. RhoA functions as a regulatory switch that confers increased endothelial cell permeability [19] and BBB disruption [20] when activated. We assessed the involvement of GXM in mediating RhoA activation in endothelial cells and found that RhoA activation by GXM is significant and time-dependent, suggesting that disruption of cytoskeletal regulation in endothelial cells may be a mechanism of virulence for *C. neoformans* to cross the BBB and establish infection in the brain. These findings complement our *in vitro* observations that GXM alters BBB permeability and contributes to increased fungal transmigration into the brain.

## RESULTS

### GXM concentrations in murine tissues 24 h post-administration

Given that we explored the impact of exogenous GXM on *C. neoformans* H99 strain infection, we first determined whether GXM disseminated systemically from the airways and could be detected in serum and brain tissue (Table 1). Therefore, C57BL/6 mice (*n* = 5) were challenged i.n. with a single GXM dose of 125 µg/mL, and the CPS levels were quantified by enzyme-linked immunosorbent assay (ELISA) 24 h later. Lung tissue GXM levels were 2.17 ± 0.09 µg/g. GXM concentrations in serum and brain tissue were 0.34 ± 0.05 µg/mL and 0.21 ± 0.13 µg/g, respectively. These measurements validated that GXM delivered into the nasal cavity can be detected in tissues other than the lungs.

**Table 1.**
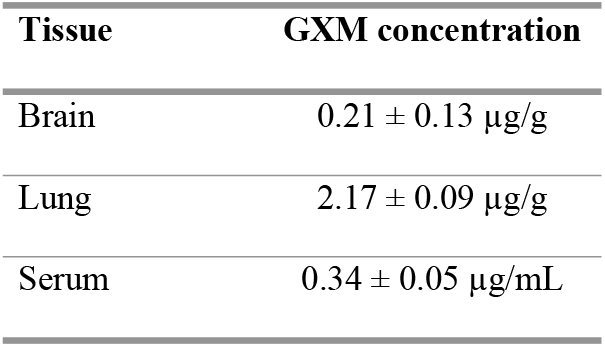
GXM levels in tissues of C57BL/6 mice (*n* = 5) 24 h post-intranasal administration.

### Pre-conditioning with GXM exacerbates murine cryptococcosis and mortality

We investigated the impact of GXM prior to C57BL/6 infection with *C. neoformans* H99 strain cells (Fig. 1). Mice were sensitized i.n. with 125 µg/mL of GXM 24 h pre-inoculation with 10^5^ *C. neoformans* yeast cells (Fig. 1A). Cryptococcosis progression was compared between these mice and rodents infected with the fungus only (untreated). i.n. administration of GXM significantly accelerated the death of *C. neoformans-*infected mice relative to control mice (*n* = 10 mice per group; *P* < 0.001; Fig. 1B). On day 16 post-infection, 100% of GXM-treated mice had died compared to 20% of untreated mice. On average, GXM-treated mice died 12-days post-infection (dpi), whereas untreated mice died 22-dpi (Fig. 1B). These findings indicate that i.n. GXM sensitization enhances mortality in mice upon pulmonary infection.

**Fig. 1.**
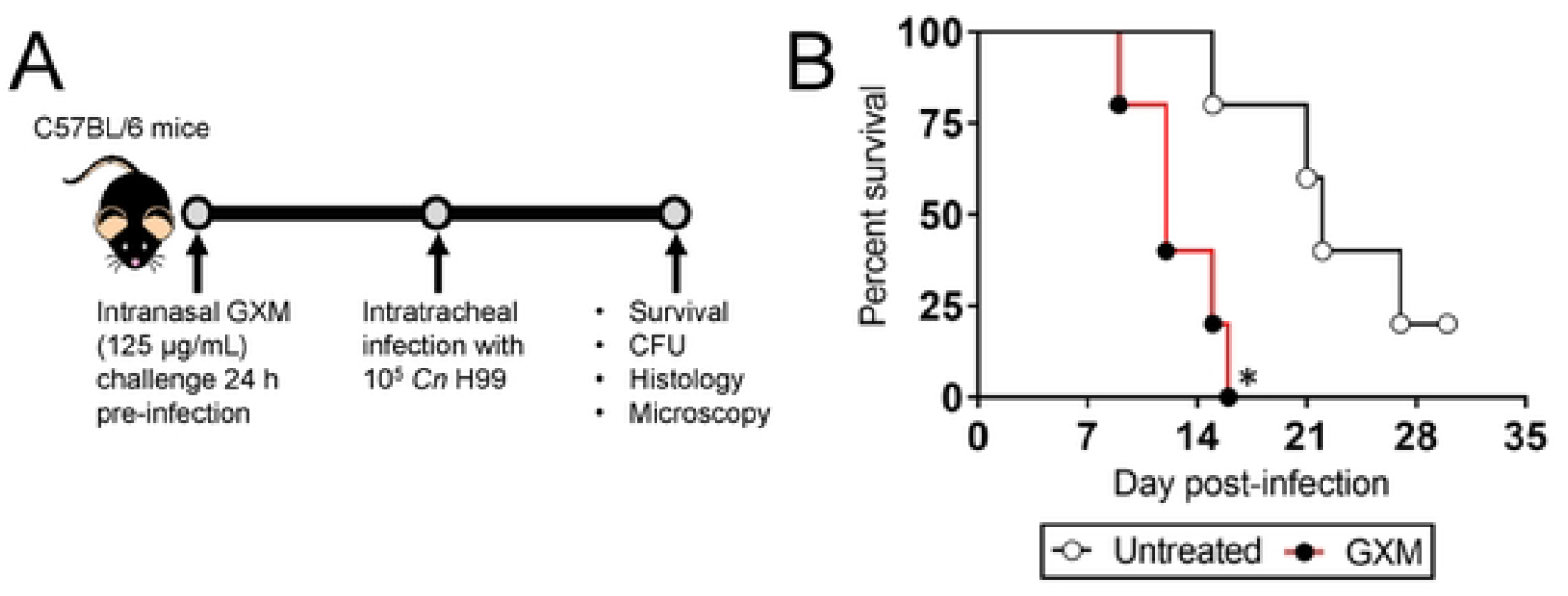
Exogenous glucuronoxylomannan (GXM) administration and *Cryptococcus neoformans* H99 strain pulmonary infection reduces survival in C57BL/6 mice. **(A)** Experimental timeline for the intranasal (i.n.) GXM challenge and fungal intratracheal infection model. Mice were sensitized with 125 µg/mL of GXM 24 h pre-inoculation with 10^5^ *C. neoformans* cells. Then, survival studies and colony forming units (CFU) determinations, histopathology, and microscopy were performed 3-and 7-days post-infection (dpi). **(B)** Survival differences of untreated (e.g., sterile phosphate-buffered saline (PBS)-sensitization) and GXM-treated C57BL/6 mice infected with 10^5^ fungi (*n* = 10 per group). *P*-value significance (* *P* < 0.05) was calculated by log rank (Mantel-Cox) analysis. This experiment was performed twice, similar results were obtained each time, and all the results combined are presented.

### Exogenous GXM enhances pulmonary and systemic fungal load

We quantified *C. neoformans* pulmonary burden in untreated and GXM-treated mice 3- and 7-dpi. Although we did not observe cryptococcal load differences between untreated [4.1 × 10^4^ colony forming units (CFU)] and GXM (4.59 × 10^4^ CFU)-challenged rodents 3-dpi (Fig. 2A), we demonstrated that the difference in pulmonary fungal burden was evident 7-dpi, with GXM-treated animals (4.66 × 10^4^ CFU) having a significantly higher burden than untreated mice (2.35 × 10^2^ CFU; *P*<0.0001; Fig. 2A). Interestingly, untreated mice infected with cryptococci showed a considerable reduction in fungal load 7-dpi (4.1 × 10^4^ CFU) relative to 3-dpi (2.35 × 10^2^ CFU). Hematoxylin and eosin (H&E) staining of pulmonary tissue removed from untreated mice 7-dpi evinced localized inflammation and dense cellular infiltration around aggregates or individual cryptococci (Fig. 2B; left upper panel; 20X). Mucine carmine staining of magnified lung tissue from untreated rodents showed *C. neoformans* cells (black arrows) of diverse sizes surrounded by inflammatory cells (red arrowheads) with limited or contained CPS production particularly around individual aggregates or individual cryptococci (Fig. 2B; left lower panel; 40X). In contrast, lungs excised from GXM-treated mice exhibited high fungal burden characterized by large numbers of cryptococci embedded in abundant amounts of CPS with little inflammation (Fig. 2B; right upper [40X] and lower [20X] panels). In addition, cryptococci was more extensively found in the blood and brain of GXM-treated mice when compared to the untreated group (Fig. 2C). These results suggest that GXM dysregulates host homeostasis and enhances disease by increasing pulmonary fungal burden, inhibiting host inflammation, facilitating the access of cryptococci to the bloodstream, and enhancing fungal dissemination to the CNS.

**Fig. 2.**
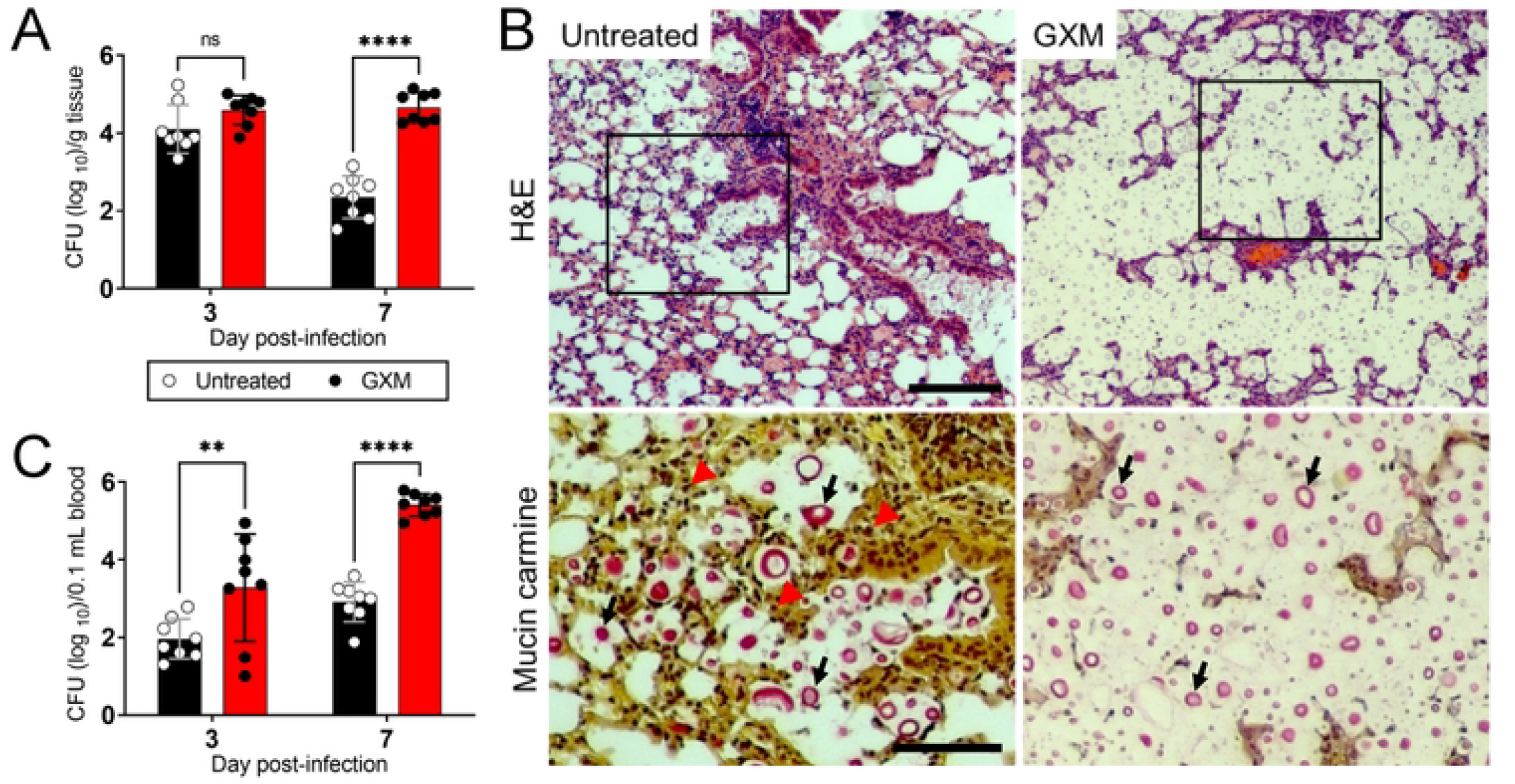
GXM exacerbates pulmonary cryptococcosis and systemic fungal load. Fungal burden in **(A)** lungs (numbers of CFU/gram (g) of tissue) and **(B)** blood (numbers of CFU/0.1 mL of blood) collected from untreated and GXM-treated mice i.t. infected with 10^5^ cryptococci (*n* = 8 per group) 3- and 7-dpi. For panels **B** and **C**, bars and error bars denote the means and standard deviations (SDs), respectively. Asterisks indicate *P-*value significance (** *P* < 0.01 and **** *P* < 0.0001) calculated using multiple student’s *t*-test analyses. ns represents not statistically significant comparisons. This experiment was performed twice, similar results were obtained each time, and all the results combined are presented. **(C)** Histological analysis of lungs removed from untreated (left panels) and GXM-treated (right panels) C57BL/6 mice infected with 10^5^ *C. neoformans* H99 strain cells. Representative H&E (host tissue morphology; upper panels; scale bar: 100 µm)- and mucin carmine (fungi; red staining; lower panels)-stained sections of the lungs are shown. Tissues were removed 7-dpi. Lower panel images are magnifications of the smaller boxes in the H&E-stained sections to better show the cryptococcal cells stained with mucin carmine (indicated with arrows; scale bar: 20 µm).

### GXM facilitates *C. neoformans* dissemination from the lungs to the CNS

We investigated the impact of exogenous GXM administration on *C. neoformans* dissemination to the brain (Fig. 3). We demonstrated that brain fungal load in GXM-treated animals (3-dpi, 2.45 × 10^2^ CFU; 7-dpi, 5.03 × 10^5^ CFU) were significantly higher than in untreated (3-dpi, 1.67 × 10^1^ CFU; 7-dpi, 1.94 × 10^1^ CFU) mice (3-dpi, *P* < 0.05; 7-dpi, *P* < 0.0001; Fig. 3A). IF staining of *C. neoformans-*infected brain tissue showed cryptococci of different size (yellow arrowheads) embedded in neuronal tissue (Fig. 3B). A tissue section of the dentate gyrus in the hippocampus of a GXM-challenged mouse exhibited a large cryptococcoma (red arrows) filled with yeasts cells (yellow arrowhead) and substantial amounts of capsular GXM released (white arrowheads) (Fig. 3C). Extensive accumulation of GXM (white arrowheads) was also observed in cortical tissue (Fig. 3D) and a large blood vessel (Fig. 3E) in the brain parenchyma of GXM-treated and *C. neoformans*-infected mice. IF staining showed a significantly higher number of lesions in the brains of GXM-treated mice (average, 7.86 ± 0.91) than those of untreated animals (2.43 ± 0.43) (*P* < 0.001) (Fig. 4A-B). The average areas of brain lesions of GXM-treated infected mice reached 241 μm^2^ ± 27.79, whereas lesions of control mice averaged 166 μm^2^ ± 75 (*P* < 0.05) (Fig. 4C). These studies demonstrate that exogenous GXM administration increases the permeability of the CNS to *C. neoformans* invasion and colonization.

**Fig. 3.**
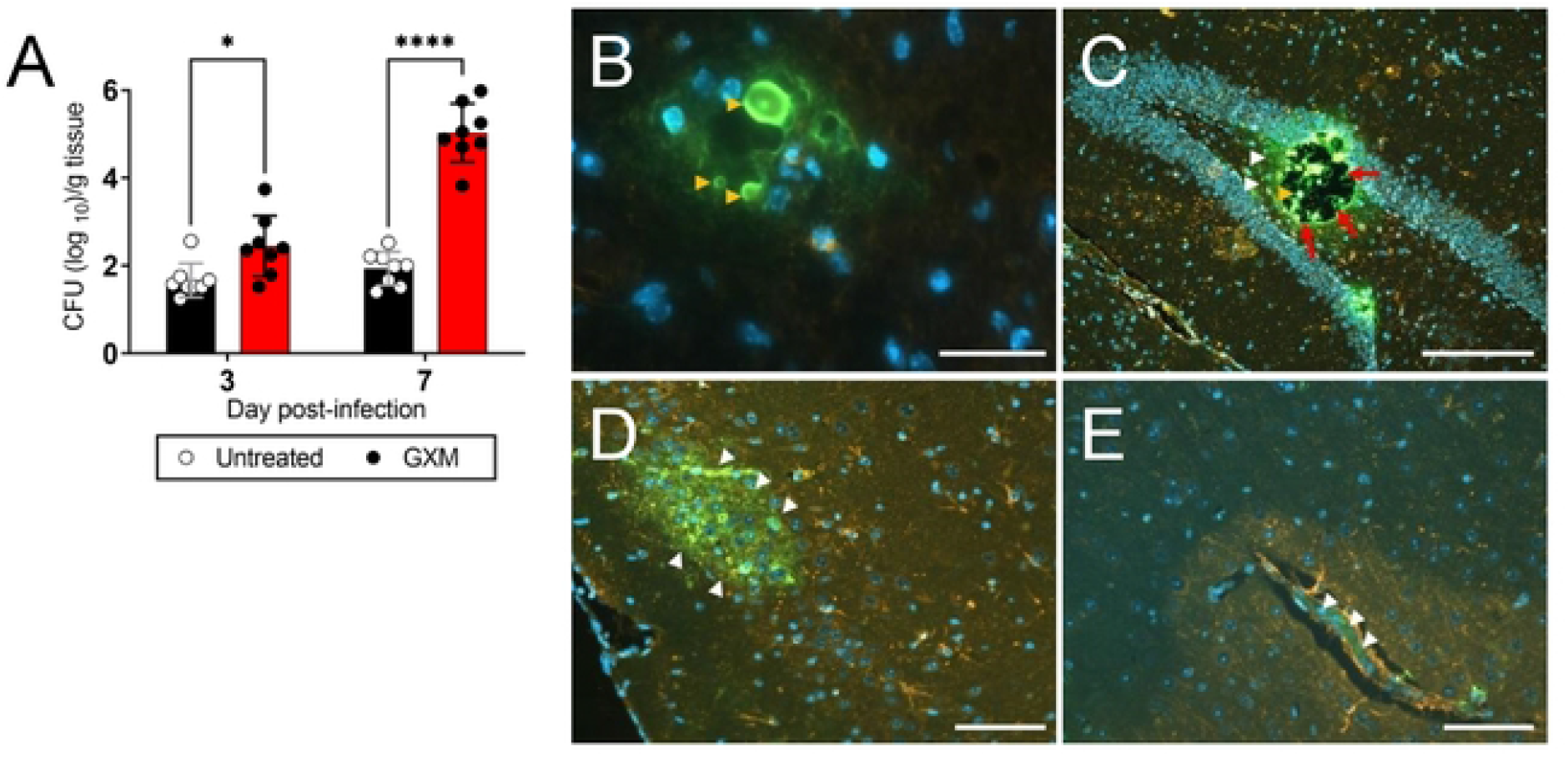
Exogenous GXM challenge enhances *C. neoformans* migration into the central nervous system (CNS) of C57BL/6 mice. **(A)** Fungal burden (numbers of CFU/gram of tissue) in brain collected from untreated and GXM-treated mice i.t. infected with 10^5^ cryptococci (*n* = 8 per group) 3- and 7-dpi. Bars and error bars denote the means and SDs, respectively. Asterisks denote *P-*value significance (* *P* < 0.05 and **** *P* < 0.0001) calculated using multiple student’s *t*-test analyses. **(B)** Confocal microscopy of cryptococcal cells (yellow arrowheads) in the brain parenchyma of GXM-treated mice. Scale bar: 20 µm. **(C)** Immunofluorescent (IF) image of a hippocampal tissue section displaying a large cryptococcoma (red arrows) filled with yeasts cells (yellow arrowhead) and abundant amounts of capsular polysaccharide released (white arrowheads). Scale bar: 150 µm. **(D)** GXM accumulation (white arrowheads) in the cortex and **(E)** blood vessel of GXM-treated and *C. neoformans* infected mice. Scale bar: 200 µm. For **B** to **E**, FITC-labeled (green) GXM-specific monoclonal antibody (mAb) 18B7 was used to label cryptococci. MAP-2 (red) and DAPI (blue) staining were used to label the cell bodies and nuclei of neurons, respectively.

**Fig. 4.**
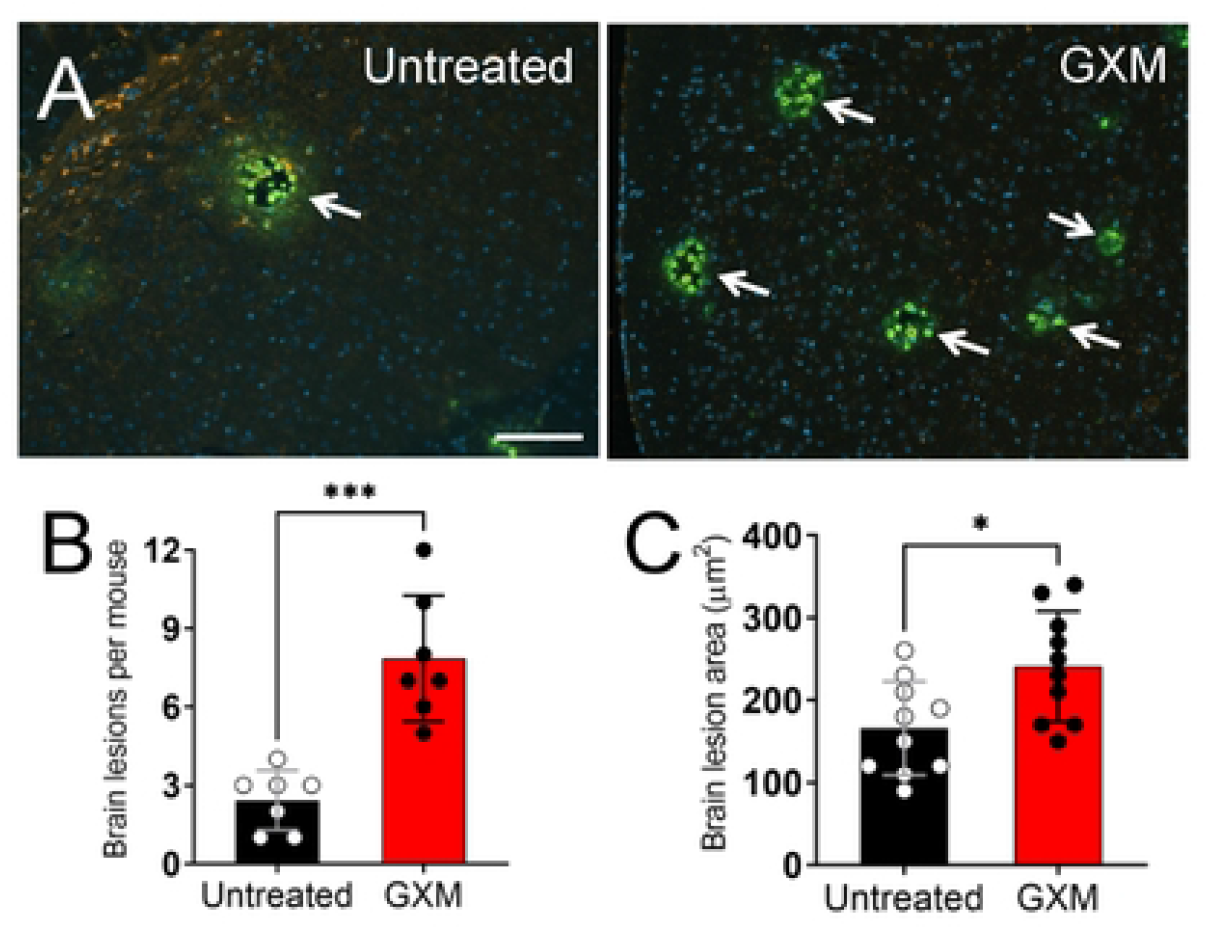
Mice sensitized with exogenous GXM exhibit more and larger brain lesions or cryptococcomas than untreated animals. **(A)** IF staining of brain lesions (cryptococcomas; arrows) caused by *C. neoformans* 7-dpi in untreated or GXM-treated animals. Fluorescein isothiocyanate (FITC)-labeled (green) capsular-specific mAb 18B7 was used to label fungal cells. MAP-2 (red) and DAPI (blue) staining were used to label the cell bodies and nuclei of neurons, respectively. Scale bar: 150 µm. **(B)** Brain lesions per mouse (*n* = 7 per group) and **(C)** lesion area (*n* = 10 per group) analyses in tissue sections of untreated and GXM-treated mice infected with *C. neoformans*. The areas of 10 brain lesions per condition were measured using NIH ImageJ software. Each symbol represents a single lesion. Bars and error bars denote the mean value and SDs, respectively. Asterisks denote *P*-value significance (* *P* < 0.05 and *** *P* < 0.001) calculated using student’s *t*-test analysis.

### GXM reduces the expression of TJs proteins in mouse brain tissue

To confirm that the CPS reached the CNS, immunofluorescence (IF) staining using GXM-specific monoclonal Ab (mAb) 18B7 demonstrated fungal polysaccharide (green) distribution (white arrowheads) throughout the brain parenchyma 24 h after administration (Fig. 5A). During pathologic conditions, TJ disruptions promote enhanced leukocyte adhesion to, and transmigration across, CNS vessels [21], resulting in BBB permeability and parenchymal leukocyte accumulation [22, 23]. Presently, studies dealing with the effects of GXM on the expression of TJs in the mouse brain are scarce. TJ proteins provide essential structural support to the BBB playing an important role in maintaining a safe neural microenvironment in the brain. Using Western blot (WB) analysis (Fig. 5B), we found that GXM administration prior to *C. neoformans* pulmonary infection alters specific TJ expression in the mouse brain. GXM doses ≥ 62.5 µg/mL decrease the expression of claudin-5 (Fig. 5C), ZO-1 (Fig. 5D), and JAM-A (Fig. 5F; *P* < 0.05) in brain tissue. Interestingly, occludin downregulation was observed in rodent brains challenged with GXM doses ≥ 15.6 µg/mL (Fig. 5E). Our data suggest that GXM-induced TJ protein reduction may be a major cause of profound brain alterations including an increase in susceptibility of the CNS to cryptococcal infection.

**Fig. 5.**
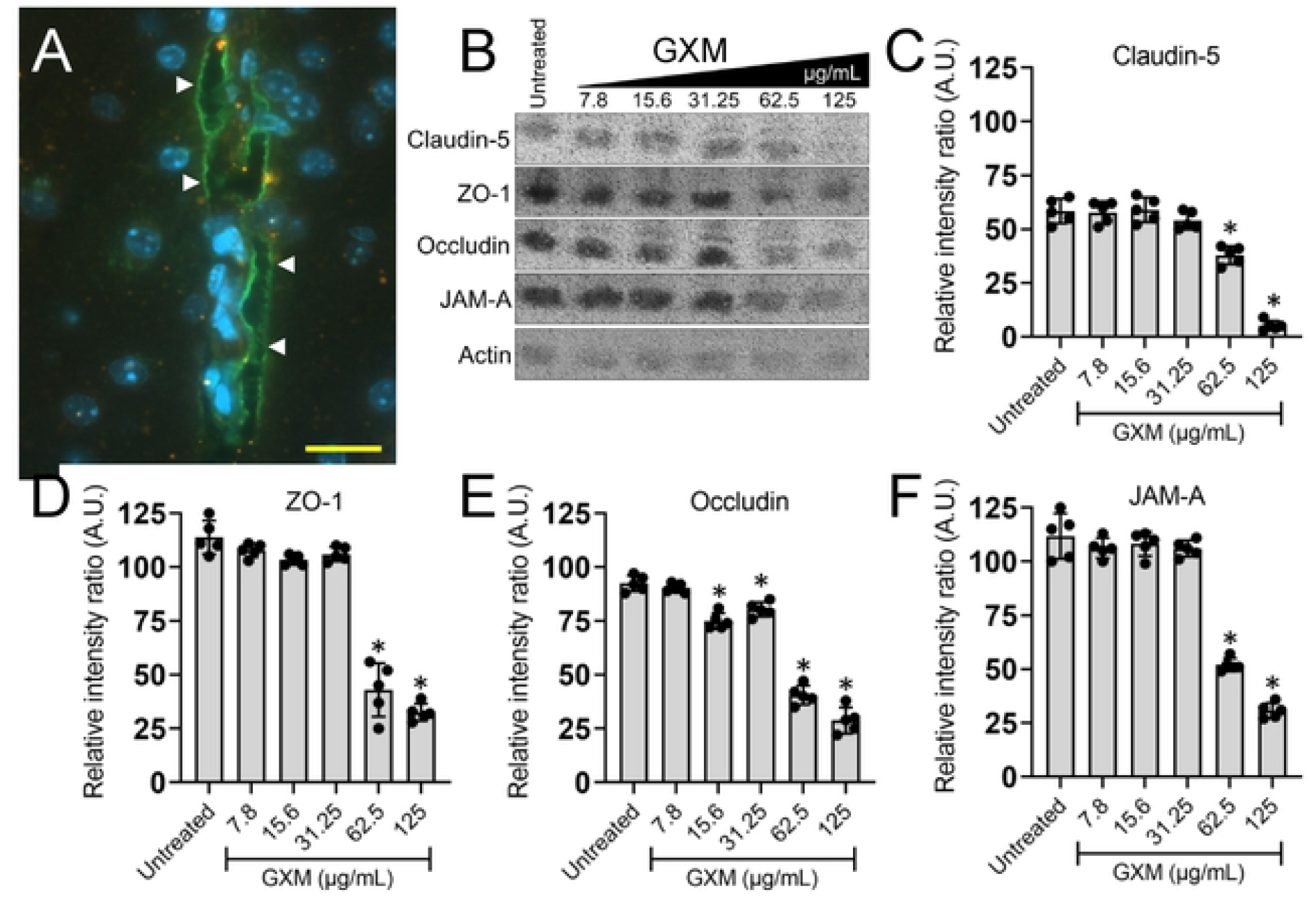
GXM challenge alters tight junction protein (TJ) expressions *in vivo*. **(A)** Confocal microscopy of cryptococcal GXM [FITC-labeled (green; arrowheads) GXM-specific mAb 18B7] distribution in the brain parenchyma (blue; nuclei of neurons) of GXM-treated mice 24 h after i.n. administration. Scale bar: 20 µm. **(B)** Western blot (WB) analyses of brain tissue from untreated (e.g., sterile PBS-sensitization) or GXM (7.8, 15.6, 31.25, 62.5, and 125 µg/mL)-challenged C57BL/6 mice were performed to compare the expression of the TJ proteins claudin-5 (23 kDa), ZO-1 (200 kDa), and occludin (59 kDa) and the adhesion protein JAM-A (32 kDa). Actin (42 kDa) was used as a housekeeping protein control. Quantitative measurements of individual band intensity in WB analyses described in panel **B** for **(C)** claudin-5, **(D)** ZO-1, **(E)** occludin, and **(F)** JAM-A, using NIH ImageJ software. Actin was used as a control to determine the relative intensity ratio (A.U. denotes arbitrary units). Bars represent the means of 5 independent gel results (black circles) and error bars indicate SDs. Asterisks denote *P*-value significance calculated using ANOVA and adjusted using the Tukey’s post-hoc analysis. \**P* < 0.05, for the reduction in the intensity of the band of TJ and adhesion molecules as compared to actin.

### GXM alters the distribution/intensity of TJs on human brain endothelial cells (HBECs)

To validate the results obtained in the murine model and to determine the impact of GXM on TJs, we analyzed the distribution of TJs (claudin-5 and occludin) in HBECs after exposure to the capsular component for 4 h using IF microscopy (Fig. 6). The distribution of claudin and occludin on HBECs is considerably reduced after incubation with 10 μg/mL of GXM (Fig. 6A). Ethylenediaminetetraacetic acid (EDTA; 10 μg/mL) was used as a positive control [24] and, similarly to GXM, substantially decreased the distribution of TJs on the surface of HBECs. Quantification of claudin-5 intensity on HBECs using the NIH ImageJ software demonstrated that GXM (*P* < 0.01) and EDTA (*P* < 0.0001) significantly reduced this TJ intensity relative to untreated control HBECs (Fig. 6B). There were no differences in the distribution of claudin-5 in HBECs incubated with EDTA or GXM. Occludin intensity was also significantly decreased in HBECs cultured with GXM (*P* < 0.05) or EDTA (*P* < 0.0001) compared to untreated cells (Fig. 6C). Similarly, EDTA-treated HBECs evinced lower occludin intensity on their surfaces than GXM-treated cells (*P* < 0.01). These experiments indicate that the distribution/intensity of TJs on HBECs is reduced after incubation with GXM.

**Fig. 6.**
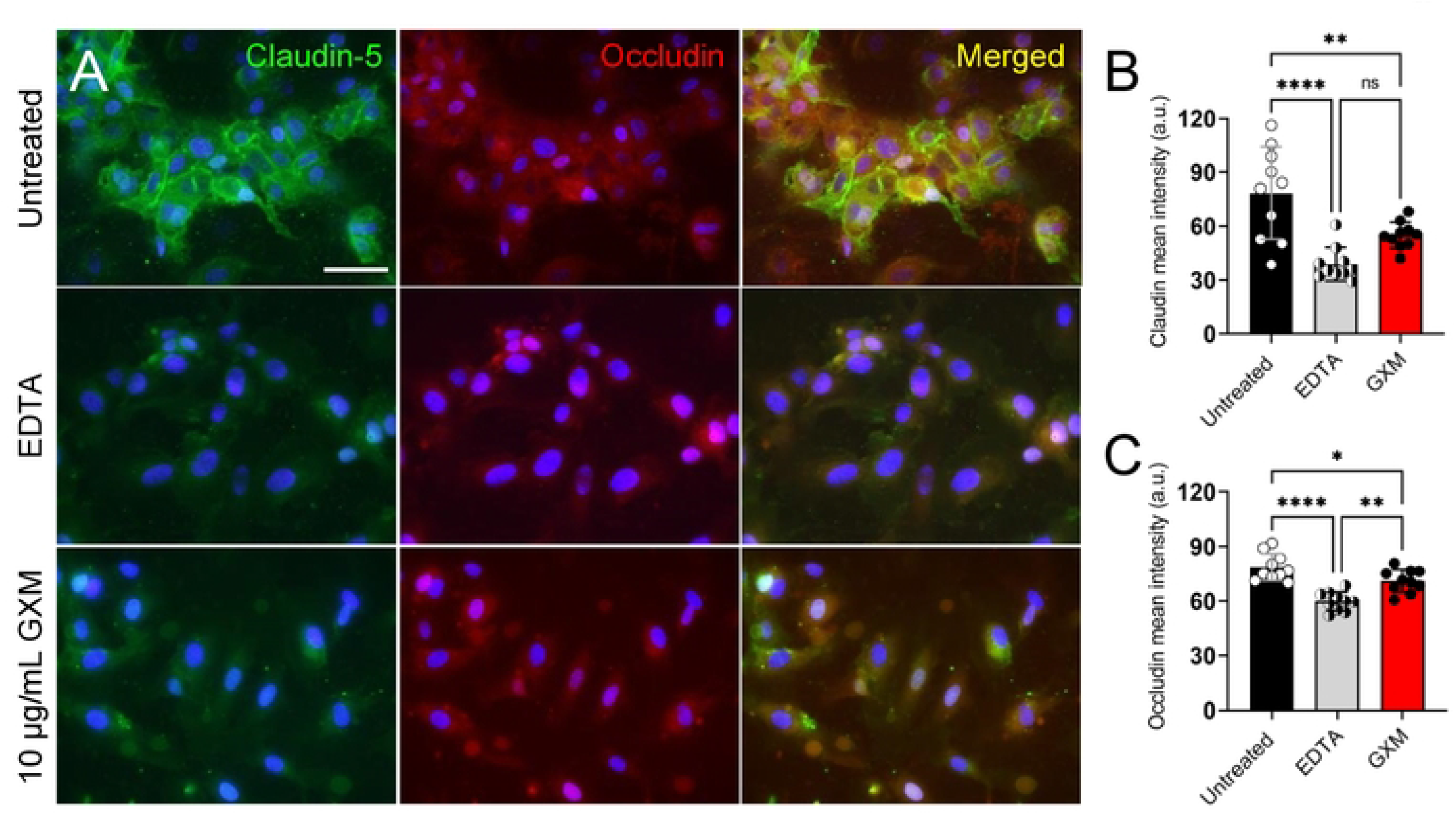
GXM alters the distribution of TJs on human brain endothelial cells (HBECs). **(A)** IF images of claudin and occludin distribution on the surface of HBECs after incubation in absence (untreated) and presence of 10 μg/mL of ethylenediaminetetraacetic acid (EDTA; positive control) or GXM for 4 h at 37 °C and 5% CO_2_. After co-incubation with EDTA or GXM, the cells were washed and incubated with claudin-FITC (green) and occludin-rhodamine (red)-specific antibodies. DAPI (blue) was used to stain the cell nuclei. Scale bar, 100 μm. **(B)** Quantification of **(B)** claudin and **(C)** occludin intensity on the surface of HBECs was performed using NIH ImageJ software. Bars and error bars denote the means and SDs, respectively. Each symbol denotes a single cell measurement (*n* = 10 cells per group). Asterisks indicate *P*-value significance (* *P* < 0.05, ** *P* < 0.01 and **** *P* < 0.0001) calculated using ANOVA and adjusted using the Tukey’s post-hoc analysis. ns represents not statistically significant comparisons.

### GXM increases the expression of RhoA cytoskeletal regulator in time-dependent manner

A previous study demonstrated that *C. neoformans* activates RhoGTPases to promote transmigration across a HBEC monolayer *in vitro*, which is the critical step for cryptococcal brain infection and development of cryptococcal meningoencephalitis (CME) [25]. To confirm that TJ integrity is disrupted by GXM, levels of both RhoA (pan) and RhoA-GTP (activated form) were measured in lysates of HBECs by WB analysis (Fig. 7). HBECs (*n* = 3) were plated in 24-well plates and grown to confluency prior to stimulation with 10 µg/mL of GXM over time (0, 15, 30, 60, 120, and 180 min; Fig. 7A). RhoA activation was induced immediately upon stimulation by GXM, reaching a peak 6-fold increase after 30 min (Fig. 7B). Interestingly, these levels dropped to near-basal levels after 60 min before decreasing below basal levels at 180 min. RhoA activation is associated with increased cell permeability and cytoskeletal fiber formation [19], and its involvement in modulating TJ integrity is of interest during CME. Actin was used as a control, exhibiting a decrease in expression over the course of the time points indicated and providing support for the role of GXM in modulating cytoskeletal properties. Our data suggest that GXM-induced RhoA activation at 30 min, followed by a prominent decrease after 60 min, may be mechanistically important for the paracellular crossing of *C. neoformans* as free yeast cells or inside of macrophages through the BBB during pathologic conditions.

**Fig. 7.**
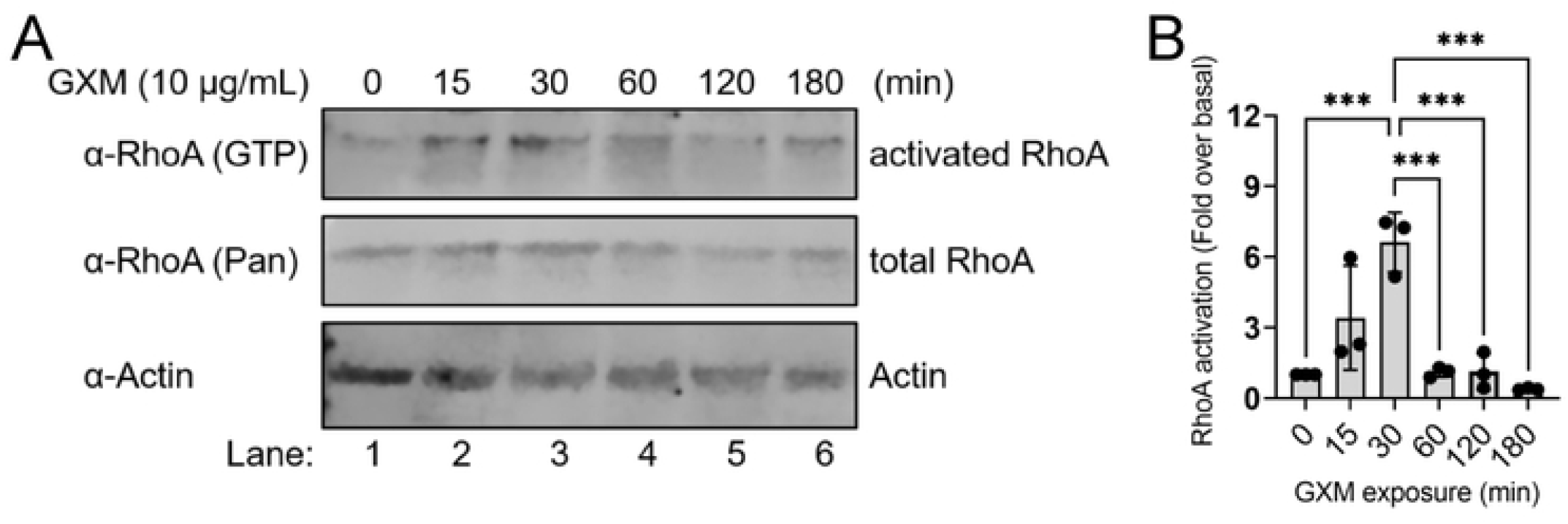
GXM augments the expression of activated RhoA in HBECs in a time-dependent manner. **(A)** WB analyses of HBEC lysates stimulated with 10 μg/mL of GXM were performed to assess the expression of activated RhoA protein (GTP-bound) over the course of 180 min. Actin was used as a protein control. Quantitative measurements of individual band intensity in WB analyses described in panel **A** for **(B)** activated RhoA using NIH ImageJ software. Bars represent the means of 3 independent gel results (black circles) and error bars indicate SDs. Asterisks denote *P*-value significance calculated using ANOVA and adjusted using the Tukey’s post-hoc analysis. \*\*\**P* < 0.001 represents a significant increase in activated RhoA after 30 min compared to the other time points.

### GXM disrupts the trans-endothelial electrical resistance (TEER) of HBECs in a blood brain barrier (BBB) model and enhances its permeability

The TEER is a widely accepted quantitative technique to measure the integrity of TJ dynamics in cell culture models of endothelial monolayers. Since GXM reduces the expression (*in vivo*) and distribution/intensity (*in vitro*) of TJs, we investigated the impact of the CPS on TEER of an *in vitro* BBB model that consisted of HBECs and pericytes (Fig. 8A-B). Pericytes are cells present at intervals along the walls of blood vessels. In the CNS, they are important for blood vessel formation, maintenance of the BBB, regulation of immune cell entry to the CNS, and control of brain blood flow [26]. Currently, there is no data available on the role of pericytes in cerebral cryptococcosis. Using this model, all the BBBs treated with GXM or EDTA demonstrated a significant drop in TEER (∼ 60% or more) 1 h post-exposure relative to the untreated BBBs (*P* < 0.05; Fig. 8B). EDTA-treated BBBs demonstrated a lower TEER percentage than BBBs incubated with 50 (*P* < 0.05) and 100 (*P* < 0.01) μg/mL of GXM. Also, BBBs cultured with 100 μg/mL of GXM exhibited lower TEER percentage than BBBs exposed to 10 μg/mL of the fungal polysaccharide (*P* < 0.01). BBBs treated with 10 μg/mL of GXM maintained ∼ 60% TEER percentage reduction after 1 h compared to the control BBBs. Notably, all the other conditions demonstrated a 100% TEER or TJ integrity reduction in BBBs after 1 h. Then, we assessed the BBB permeability to streptavidin-horseradish peroxidase (HRP) after 1 h treatment with GXM or EDTA (Fig. 8C-D). All the GXM (10 μg/mL, *P* < 0.05; 50 μg/mL, *P* < 0.01; 100 μg/mL, *P* < 0.001)-or EDTA (*P* < 0.001)-treated HBECs exhibited significantly more permeability than the untreated cells (Fig. 8D). HBECs cultured with concentrations ≥ 50 μg/mL of GXM or 10 μg/mL of EDTA displayed similar increased in barrier permeability to streptavidin-HRP. HBECs incubated with 10 μg/mL of GXM demonstrated considerably lower barrier permeability than EDTA-treated (*P* < 0.05) cells, respectively. Our experiments demonstrated that GXM disturbs the TEER and integrity of HBEC-closed interactions in a BBB *in vitro* model.

**Fig. 8.**
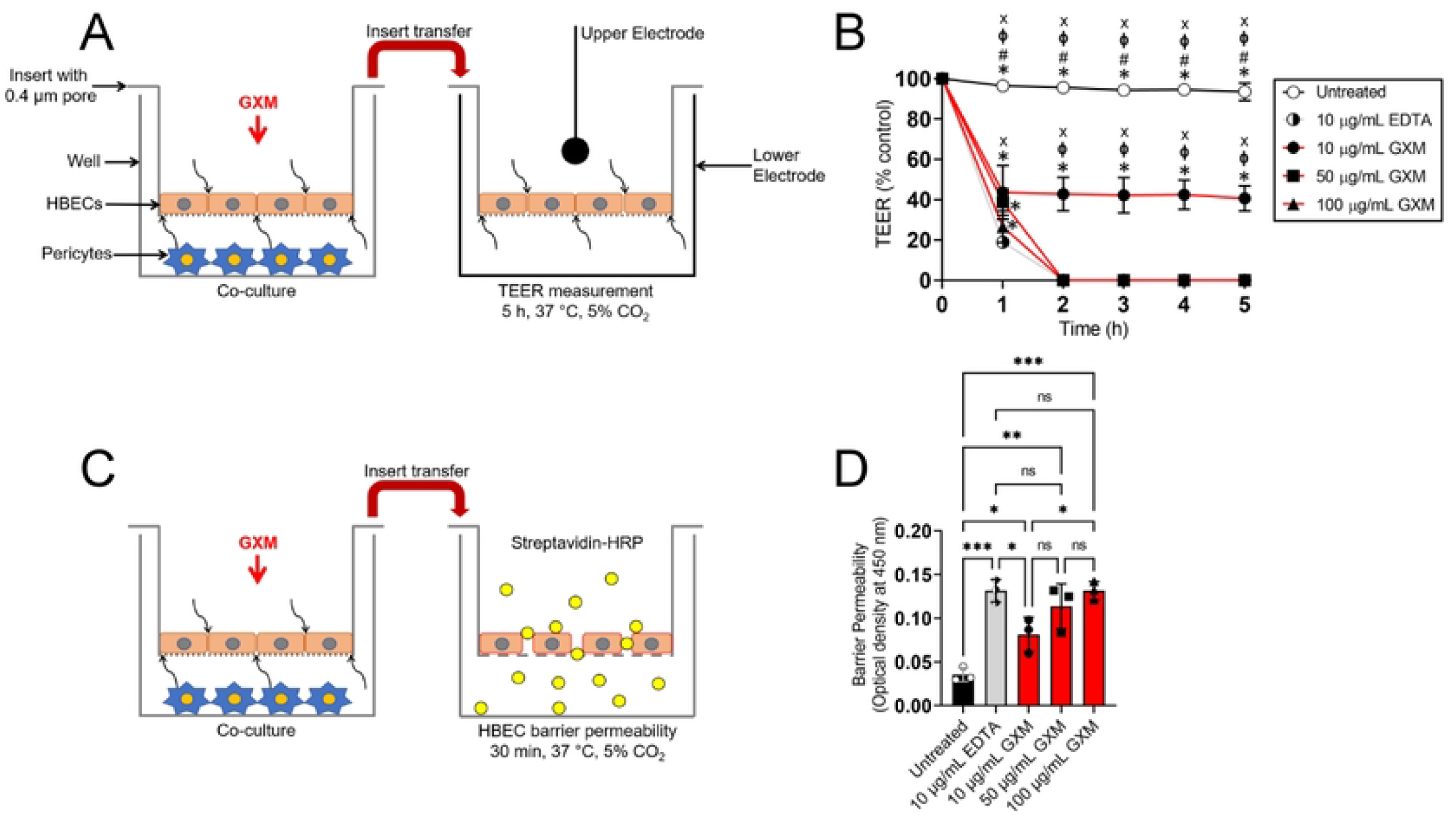
GXM decreases the trans-endothelial electrical resistance (TEER) and increases the permeability of HBECs in a blood brain barrier (BBB) model. Graphic representation of the HBECs and pericytes BBB model used for the **(A)** TEER measurements and **(C)** HBEC barrier permeability assay. HBECs and pericytes were grown separately (HBECs, 0.4 μm pore insert; pericytes, bottom of a well) and co-incubated using a microtiter transwell system that permits chemotactic exchange through the supernatant. **(B)** Relative TEER of HBECs incubated with GXM (10, 50, or 100 μg/mL) for 5 h at 37 °C and 5% CO_2_. HBECs incubated alone or with EDTA (10 μg/mL) were used as negative and positive controls, respectively. Time points are the averages of the results for three different TEER measurements (*n* = 3 wells per group per experiment), and error bars denote SDs. Symbols (*, #, ϕ, and x) indicate *P* value significance (*P* < 0.05) calculated using ANOVA and adjusted using the Tukey’s post-hoc analysis. *, #, ϕ, and x indicate significantly higher TEER than in the 10 μg/mL of EDTA-, 10 μg/mL of GXM-, 50 μg/mL of GXM-, and 100 μg/mL of GXM-treated groups, respectively. **(D)** Relative HBEC barrier permeability to streptavidin-horseradish peroxidase (HRP) after incubation with GXM (10, 50, or 100 μg/mL) for 30 min at 37 °C and 5% CO_2_. HBECs incubated alone or with EDTA (10 μg/mL) were used as negative and positive controls, respectively. Bars and error bars denote the means and SDs, respectively. Each symbol denotes a single well measurement (*n* = 3 wells per group). Asterisks indicate *P*-value significance (* *P* < 0.05, ** *P* < 0.01 and *** *P* < 0.001) calculated using ANOVA and adjusted using the Tukey’s post-hoc analysis. ns represents not statistically significant comparisons. These experiments were performed thrice, similar results were obtained each time, and all the results combined are presented.

### GXM facilitates the movement of *C. neoformans* through HBECs

Given that GXM inhibits the expression of TJs and alters the TEER in HBECs, we investigated the permeability of HBECs to *C. neoformans* strain H99 cell transmigration after pre-incubation with 10 μg/mL of GXM or EDTA for 1 h (Fig. 9A). GXM-(*P* < 0.01) and EDTA-(*P* < 0.0001) treated HBECs demonstrated significant passage of cryptococci compared to untreated cells (Fig. 9B). EDTA had the highest cryptococcal BBB transmigration, followed by GXM and untreated HBECs, respectively. Our studies indicate that GXM promotes HBEC disruption and *C. neoformans* penetration.

**Fig. 9.**
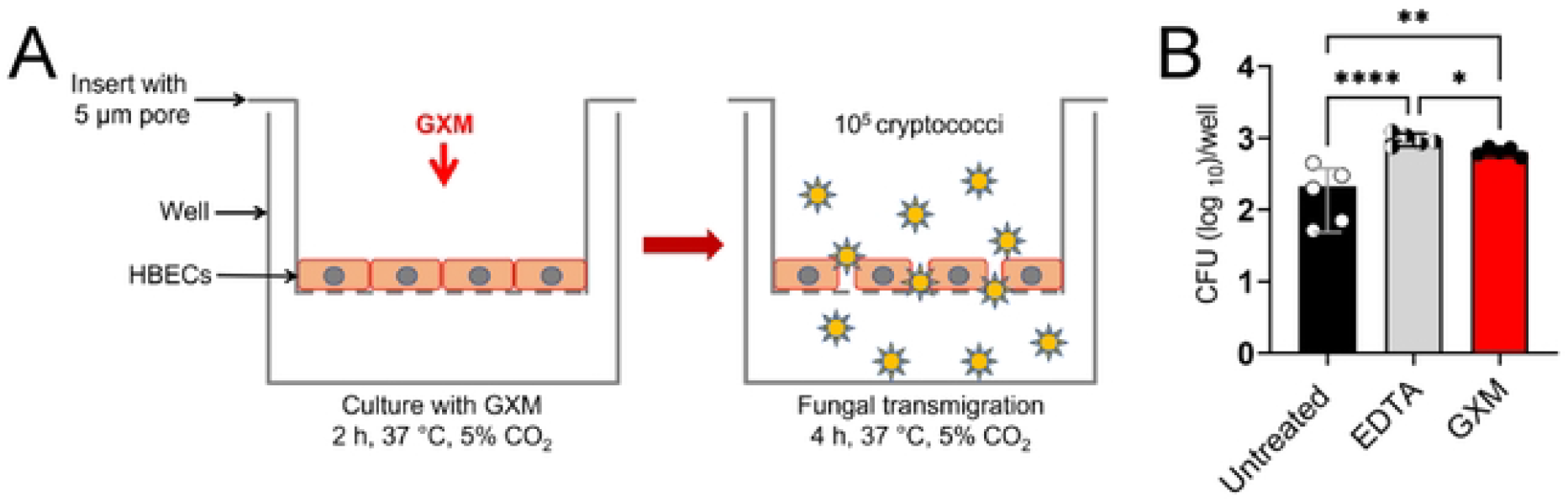
GXM increases the transmigration of *C. neoformans* through HBECs. **(A)** HBECs were cultured for 2 h with 10 μg/mL of GXM in a microtiter transwell system (5 μm pore insert). Then, 10^5^ cryptococci were added to the well and fungal transmigration through the HBECs was determined for 4 h using CFU. **(B)** CFU determinations after fungal transmigration are shown. HBECs incubated alone or with EDTA (10 μg/mL) were used as negative and positive controls, respectively. Bars and error bars denote the means and SDs, respectively. Each symbol denotes CFU determinations from a single well (*n* = 5 wells per group). Asterisks indicate *P*-value significance (* *P* < 0.05, ** *P* < 0.01 and **** *P* < 0.0001) calculated using ANOVA and adjusted using the Tukey’s post-hoc analysis. ns represents not statistically significant comparisons.

### GXM stimulates vasodilation of blood vessels via endothelium-dependent mechanisms

To understand the influence of *C. neoformans* GXM on endothelial cells of blood vessels, mouse superior mesenteric arteries (SMA) were isolated, cultured in absence or presence of 25 µg/mL of GXM *ex vivo* for 40 min, and evaluated for vascular reactivity using a wire myograph in presence of acetylcholine (ACh; to test for endothelium-dependent vasodilation) and phenylephrine (PE; to test for vasoconstriction) (Fig. 10). SMA incubated with the CPS were less responsive to PE and decreased their contraction relative to untreated arteries (*n* = 5 per group; Fig. 10A). However, arteries incubated with 25 µg/mL of GXM were substantially more sensitive to ACh and significantly increased their vasodilation compared to untreated blood vessels (Fig. 10A). We corroborated that GXM-treated SMA (*n* = 3) were significantly more sensitive to ACh than untreated arteries (*n* = 5; *P* < 0.05; Fig. 10B). We concluded that GXM promotes blood vessel relaxation by endothelium-dependent mechanisms, providing proof of principle evidence that the cryptococcal CPS alters the integrity of the vasculature, enabling systemic dissemination from the respiratory system to the CNS.

**Fig. 10.**
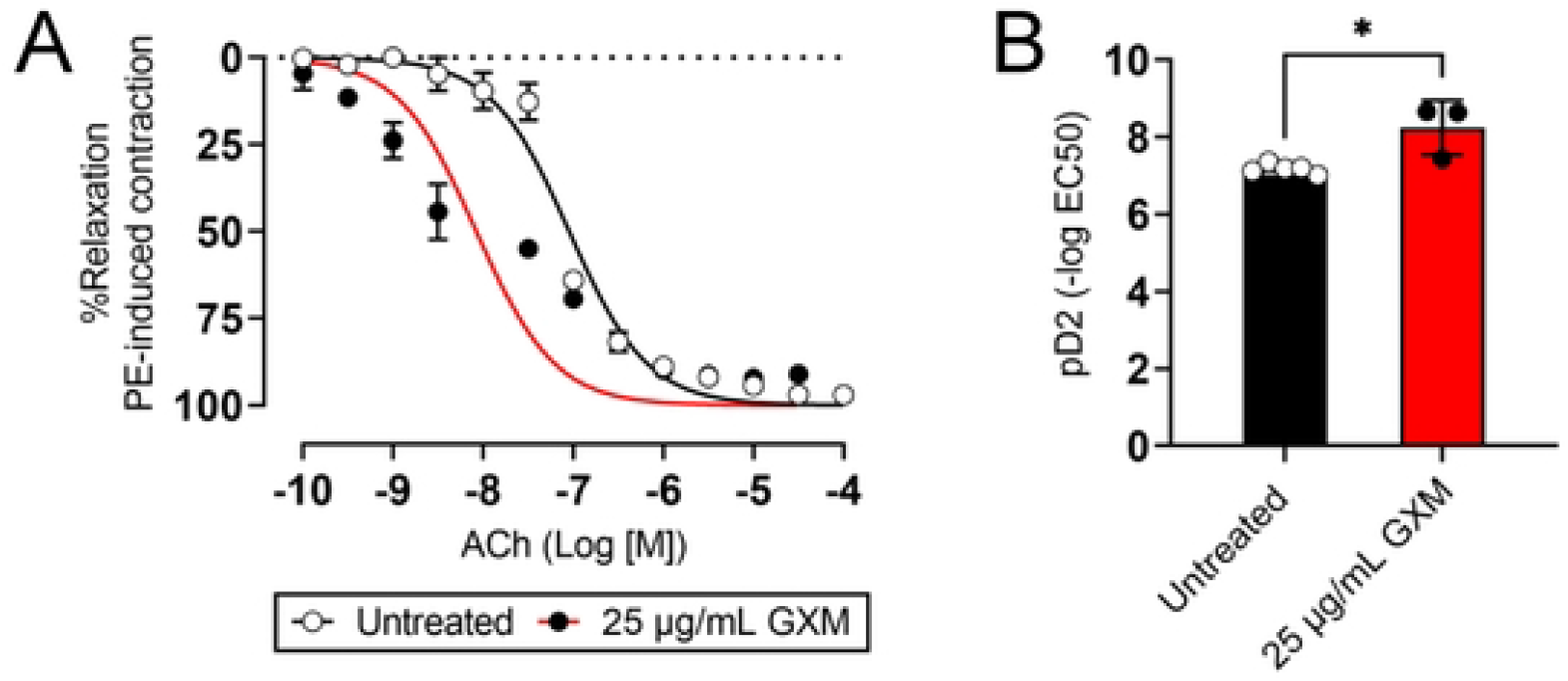
GXM causes vasodilation of mouse blood vessels after *ex vivo* treatment. **(A)** Murine superior mesenteric arteries (SMA) incubated in absence (untreated) or presence of 25 µg/mL GXM were evaluated in response to phenylephrine (PE; test for vasoconstriction) or acetylcholine (ACh; test for endothelium-dependent vasodilation). Cumulative concentration-response curves are shown. Each point represents the average of 5 individual SMA measurements. Concentration– response curve was log-transformed, normalized to percent maximal response, and fitted using a nonlinear regression. *P* values of <0.05 were considered significant. **(B)** pD2 for ACh curve. Each symbol (*n* = 5, untreated; *n* = 3, GXM-treated SMA rings) represents a single SMA ring. Bars and error bars denote the mean value and SDs, respectively. Asterisk denotes *P*-value significance (* *P* < 0.05) calculated using student’s *t*-test analysis.

## DISCUSSION

*C. neoformans* capsular GXM is released into tissues during infection where it accumulates and causes a range of adverse immunological defects [17]. Individuals with CME frequently suffer from the detrimental effects associated with increased intracranial pressure such as edema, hydrocephalus, and subarachnoid space hemorrhage, which may be related in part to GXM-mediated effects. The concentration of GXM in CSF can reach 1 mg/mL [27] and post-mortem pathological examinations of brains from patients with CME indicate that this polysaccharide is extensively deposited in tissue [17]. We evaluated the role of exogenous GXM on the development of *C. neoformans* pulmonary infection by i.n. challenging C57BL/6 mice. However, we first determined the extent of exogenous GXM dissemination post-inoculation using ELISA. GXM was detected in lung and serum in comparable concentrations to other similar studies in rodents [14, 28]. While *C. neoformans* capsular GXM is cleared from the bloodstream within days, it can be retained in organ tissues for weeks, especially those of the liver and spleen [29]. Surprisingly, we also detected low levels of GXM in brain tissue, validating that this delivery was appropriate to study the impact of the CPS on the development of cryptococcosis.

We investigated the impact of exogenous GXM on the development of *C. neoformans* infection and demonstrated that CPS accelerates cryptococcosis in C57BL/6 mice. Pulmonary cryptococcal infection of GXM-challenged rodents was characterized by substantial colonization of the epithelial tissue, extensive CPS accumulation surrounding cryptococci in a biofilm-like arrangement, and minimal inflammation. It is possible that prior priming of the respiratory tissue with GXM to cryptococcal infection enhances the ability of the fungus to adhere and colonize the airway epithelium. Although *cap67*, a strain with defective capsular production, has shown higher adhesion to human epithelial-like cells than clinical encapsulated strains [30], phospholipase B [31], mannoprotein MP84 [32], and heat-shock protein 70 [33] have been shown to influence the attachment of *C. neoformans* to pulmonary-like cells. i.n. challenge of mice with GXM may even create a competitive advantage over other possible ligands involved in the association of the carbohydrate with surface receptors on epithelial cells [33]. Moreover, active production and release of CPS by *C. neoformans* are required for biofilm formation [34], which involves a robust attachment to the surface substrate. Substantial accumulation of CPS around cryptococci in GXM-sensitized animals demonstrates a biofilm-like structure in lung tissue that prevents antifungal immune responses such as cellular migration, phagocytosis, and clearance, providing evidence for the involvement of GXM in the progression of *C. neoformans* invasion.

Given that *C. neoformans* infection was more difficult to control in the lungs of GXM-challenged mice, these animals also evinced higher hematogenous fungal load than the untreated rodents. It is conceivable that the slight immune response mediated the collapse of the lung tissue and vasodilation of the blood vessels, allowing substantial cryptococcal cell proliferation and extensive production/release of the CPS [35]. In this regard, portions (M2 motif) of the *C. neoformans* capsular GXM have been shown to elicit non-protective Abs, which may affect phagocytic cell responses in the lungs and result in the inability of the host to control the infection [36]. Mice lacking claudin-18, a TJ protein highly expressed in airway tissues, exhibited increased susceptibility to *C. deneoformans* infection through massive multiplication of yeast cells with poor granulomatous responses, reduced production of interferon-γ, and acidification of the alveolar space despite increased presence of immune cells such as CD4^+^ T cells [37]. In addition, GXM has detrimental effects on endothelial cells associated with blood vessels including alterations of adhesion molecules important for the migration of leukocytes to fight off the infection [38, 39] and vascular vasodilation [40] following cryptococcal infection, which increases vessel tension and promotes fungal dissemination [40].

GXM sensitization prior to infection increased *C. neoformans* load in the CNS. There was approximately a 3-fold CFU count difference between brains excised from untreated and GXM-challenged mice 7-dpi. Since the exogenous GXM was administered to the animals i.n., this capsular component deposited mostly in the lungs and reached serum in low levels [14]. Even though GXM injection into the bloodstream does not reach to the CNS due to the unidirectional movement of the CPS across the BBB from the CSF to serum [28, 41], the ELISA and IF data produced in this investigation demonstrated that i.n. delivery of GXM reaches the CNS likely through the nasal cavity. A recent study showed that i.n. *C. neoformans* infection was found in the upper respiratory tract and fungal brain invasion occurred quickly in ≤ 3 h [42]. GXM-treated mice showed higher number of cryptococcomas and larger cryptococcal brain lesions than those found in untreated brains. Likewise, GXM localized in blood vessels in brain parenchyma, and its deposition may have serious implications in the maintenance of the host BBB integrity. Vascular GXM accumulation and fungal occlusion have been described as responsible of infarction and hemorrhagic dissemination in CME patients [43, 44] and animal models [40], respectively. Notably, the cortex and hippocampus of GXM-challenged mice were the most affected regions of the brain, suggesting that these animals may also show behavioral and cognitive impairment. In fact, altered mental status in HIV^+^ patients with CME is associated with high mortality rates [45, 46]. Although this is outside of the scope of this study, future investigations are needed to understand the impact of cryptococcal infection and GXM on behavior and cognition.

We found that i.n. GXM instillation (concentrations: 15.6-125 μg/mL) alone disrupted TJ and adhesion proteins in rodents, validating the importance of this polysaccharide in destabilizing the BBB and promoting traversal passage of *C. neoformans* into the CNS. TJ and adhesion proteins are important in the molecular architecture of the cell and *C. neoformans* interactions with BBB endothelial cells mediate alterations in the cytoskeleton of the mammalian cells [47]. GXM has been shown to have a profound impact on HBEC TJs. Claudin-5, ZO-1, and JAM-A were downregulated at concentrations ≥ 62.5 μg/mL, whereas occludin seemed to be more susceptible to GXM and was reduced at concentrations ≥ 15.6 μg/mL. WB analyses in murine brain extracts 24 h after GXM sensitization and IF in HBECs cultured with the CPS for 4 h demonstrated a significant reduction in the expression/distribution of claudin-5 and occludin. These integral proteins are located in the apical region of the cell and act as gatekeepers for BBB paracellular passage. Our results are consistent with Chen and colleagues, who demonstrated that occludin dysregulation is associated with weakened endothelial cell interactions in the BBB [47].

It is also significant to mention that TJs impact structural changes in the cell that can be important in the transcellular passage of free cells, as well as those being transported inside of macrophages. Our findings of GXM-induced RhoA activation substantiates the impact of this CPS on the BBB structural integrity during *C. neoformans* invasion. Although the involvement of RhoGTPases in the translocation of *C. neoformans* from the blood capillaries to the brain parenchyma has been previously suggested [25], we demonstrated that GXM may regulate fungal transmigration by causing rapid structural changes in the BBB including weakening of TJ and disruption of HBEC interactions (Fig. 11). GXM stimulates activation of RhoA, which activates Rho Activated Kinase (ROCK) [47]. ROCK inhibits the myosin light chain phosphatase (MLCP), leading to actin stress fiber formation through enhancement of MLC phosphorylation [48]. Actin stress fibers contribute to the internalization and lysosomal degradation of claudin and occludin, disrupting TJs between brain microvascular endothelial cells [49]. These observations are significant because further studies can be focused on strengthening TJ interactions between neighboring HBECs in individuals with pulmonary cryptococcosis and help the development of novel strategies to prevent CME and its associated morbidity. Furthermore, investigating the role of GXM as a modulator of TJ and BBB integrity may be of interest for future studies. For instance, the receptor on the surface of HBECs that recognizes GXM and triggers the RhoA signaling pathway still needs to be identified.

**Fig. 11.**
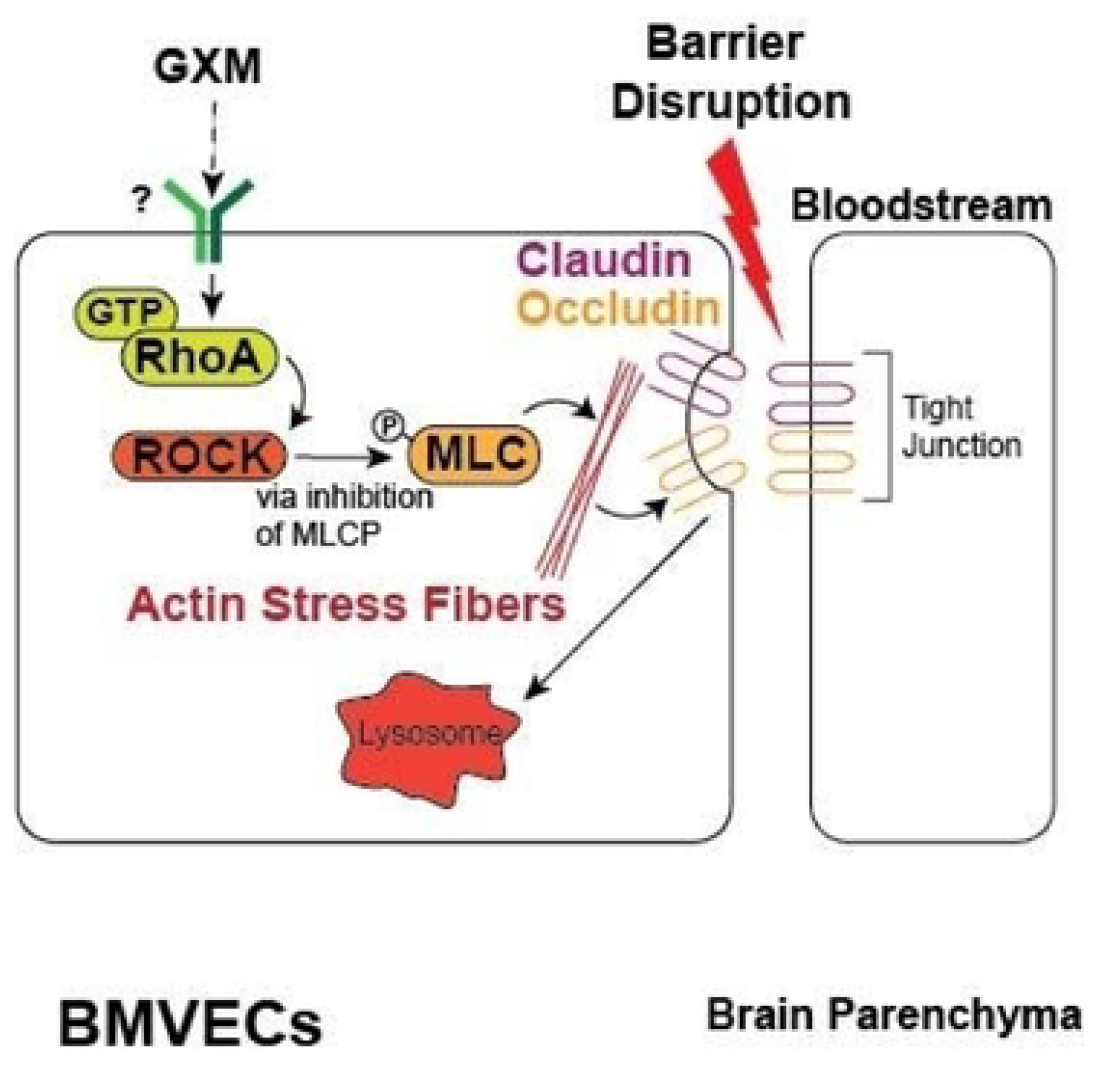
Model of GXM-mediated disruption of the BBB during cryptococcal infection. GXM stimulates activation of RhoA, which activates Rho Activated Kinase (ROCK). ROCK inhibits the myosin light chain phosphatase (MLCP), leading to actin stress fiber formation through enhancement of MLC phosphorylation. Actin stress fibers contribute to the internalization and lysosomal degradation of claudin and occludin, disrupting TJs between brain microvascular endothelial cells.

Due to the difficult nature of investigating the interaction of *C. neoformans* with the neurovascular unit *in vivo*, the use of microtiter transwell systems to mimic the BBB has proliferated in the past two decades [8, 47, 50-54]. Since we demonstrated that GXM alters the distribution of TJs on HBECs, we assessed the impact of GXM on the TEER of HBEC monolayers grown in transwells and co-incubated with pericytes separated by a membrane that allows the exchange of secreted factors by the cells in co-culture. GXM quickly reduced the TEER across the HBEC monolayer in presence of pericytes, suggesting disruptions in TJs and the integrity of the HBEC neighboring interactions. Little is known on the role of pericytes on *C. neoformans* BBB transmigration *in vivo*. However, a deficiency of pericytes in the CNS can cause increased BBB permeability due to their ability to cover endothelial cells that line the capillaries [55]. Studies to elucidate the basic functions of pericytes in *C. neoformans* transmigration require such pericyte-endothelial cell communication for a more comprehensive understanding of cerebral cryptococcosis.

Lastly, we demonstrated GXM-treated HBECs grown in transwells displayed high permeability and passage to cryptococci. It is plausible that both transmigration processes, trans-and para-cytosis, occurred in this *in vitro* system. GXM exposure to HBECs resulted in a significant reduction of TJ distribution on their surface and decreased TEER, indicating that fungal crossing between endothelial cells was taking place. We did not examine HBEC transcellular passage, but this mechanism of BBB penetration and CNS invasion has been extensively described in free cryptococci [8, 47] and cryptococci trolleying inside phagocytic cells [51]. Though the specific mechanism of *C. neoformans* brain invasion involved in this process is not yet confirmed, it may be sensible to embrace the idea that all the transmigration processes described, including the Trojan horse, simultaneously and dynamically occur during infection.

In conclusion, we demonstrated that exogenous GXM administration exacerbates lung colonization, systemic dissemination, and CNS invasion. GXM has detrimental defects on TJs facilitating its translocation from the lungs to the CNS suggesting the importance of developing therapeutics to strengthen cell-to-cell interactions, particularly in the early stages of the disease. Our findings have significant implications in understanding the different mechanisms used by the fungus to cross the BBB and colonize the brain resulting in its main manifestation, CME, which has high mortality. Additional investigations are necessary to establish the molecular mechanisms by which GXM impairs TJ function and impacts cryptococcal systemic disease.

## MATERIALS AND METHODS

### C. neoformans

*C. neoformans* strain H99 (serotype A) was isolated and kindly provided by John Perfect at Duke University. Yeasts were grown in Sabouraud dextrose broth (pH 5.6; BD Difco) for 24 h at 30°C in an orbital shaker (Thermo Fisher; TF) set at 150 rpm (to early stationary phase).

### GXM isolation

GXM was isolated from *C. neoformans* strain H99 using the hexadecyltrimethyl ammonium bromide (CTAB) method as previously described [56] with a few modifications. Briefly, fungal cells (10^9^) were inoculated into 1,000-mL Erlenmeyer flasks containing 400 mL of minimal medium composed of glucose (15 mM), MgSO_4_ (10 mM), KH_2_PO_4_ (29.4 mM), glycine (13 mM), and thiamine-HCl (3 μM), pH 5.5. Fungal cells were cultivated for 3 days at 30°C with shaking and separated from culture supernatants by centrifugation at 4,000 x g (15 min, 4°C). The supernatant fluids were collected and centrifuged again at 15,000 x g (15 min, 4°C) to remove smaller debris. The pellets were discarded, and the resulting supernatant was concentrated approximately 20-fold using an Amicon (Millipore) ultrafiltration cell (with a cutoff of 100 kDa and a total capacity of 200 mL) with stirring and Biomax polyethersulfone ultrafiltration discs (63.5 mm). A nitrogen (N2) stream was used as the pressure gas. After the supernatant was concentrated, a thick, translucent film was observed in close association with the ultrafiltration disc and was covered by a concentrated fluid phase. The fluid phase was discarded, and the viscous layer was collected with a cell scraper for storage at room temperature (RT). Fractions that were passed through the 100-kDa filtration discs were filtered through 10-kDa membranes, resulting again in film formation. We heat inactivated (100°C for 15 min) proteases in our GXM preparation. Additionally, each preparation was treated with protease inhibitor cocktail (37°C for 2 h). Each preparation was also tested for contamination with bacterial lipopolysaccharide using the Limulus amoebocyte lysate assay. For polysaccharide quantification, a capture ELISA [57], the carbazole reaction for hexuronic acid [58], and the method for hexose detection described by Dubois et al. [59] were used.

### *In vivo* GXM sensitization and pulmonary model of *C. neoformans* infection

Male and female C57BL/6 mice (10-12 weeks old; Charles Rivers) were anesthetized using the ketamine (100 mg/kg)-xylazine (10 mg/kg) cocktail (Covetrus) and i.n. challenged with a 50 µL solution of phosphate-buffered saline (PBS; untreated) or GXM (125 µg/mL) in PBS 24 h prior to cryptococcal pulmonary infection. Then, animals were anesthetized using the ketamine-xylazine cocktail and a vertical 5-mm incision was made in the skin of the ventral neck just right of midline. The trachea was identified and injected with a 100 μL suspension containing 10^5^ *C. neoformans* strain H99 yeast cells using a 26-gauge syringe. The anterior neck site was closed with surgical glue and 1% topical chlorhexidine solution was applied over the closed incision. A group of mice was monitored for survival studies. A separate group of GXM challenged/infected mice were bled from the facial vein (0.1 mL blood collected), euthanized in a chamber with 30-70% CO_2_ flow, and brain/lungs excised for CFU count determinations and histological processing. All animal studies were conducted according to the experimental practices and standards approved by the Institutional Animal Care and Use Committee at the NYIT College of Osteopathic Medicine (Protocol #: 2016-LRM-01). This study was carried out in compliance with the ARRIVE guidelines [60].

### CFU determinations

Lung and brain tissues were excised from euthanized mice 3- and 7-dpi. The right lobe of lung and the right hemisphere of the brain were homogenized in 5 mL of sterile PBS, serially diluted, a 100 µL suspension was plated on Sabouraud dextrose agar (BD Difco) and incubated at 30°C for 48 h. Quantification of viable yeast cells from untreated and GXM-challenged animals were determined by CFU counting.

### Histological examinations

#### a. Lungs

Lung tissue was excised from euthanized mice 7-dpi and fixed in 10% formalin for 24 h, processed, and embedded in paraffin. Four-micrometer vertical sections were cut and then fixed to glass slides and subjected either to hematoxylin and eosin (H&E) stain or to mucin carmine stain, to examine host tissue and fungal morphology, respectively. Microscopic examinations of tissues were performed by light microscopy with a Leica DMi8 inverted microscope (20 and 40X objectives), and photographed with a Leica DFC7000 T digital camera using the Leica software platform LAS X.

#### b. Brain

*C. neoformans* cells and CPS released in tissue were stained using GXM-specific monoclonal antibody (mAb) 18B7. Slides were blocked, and mAb 18B7 (2 µg/ml) was added for 1 h at 37°C. After the slides were washed, fluorescein isothiocyanate (FITC)-conjugated goat anti-mouse (GAM) Ab [1:250; 1% bovine serum albumin, (BSA)] was applied for 1 h at RT. Neurons in tissue sections were stained with DAPI (4′,6-diamidino-2-phenylindole) and microtubule-associated protein-2 (MAP-2) as described previously [61]. Microscopic examinations of brain sections were performed with a fully motorized Carl Zeiss Axio Observer Z1 confocal microscope. Confocal images of blue, green, and red fluorescence were conceived simultaneously using a multichannel mode. *Z*-stack images and measurements were corrected utilizing Zen software in deconvolution mode.

### Determination of GXM levels

*C. neoformans* capsular GXM in organs and serum were measured by capture ELISA as described [62] and modified accordingly for this study. Organ tissue (0.2 g) was placed in 2 mL of PBS, homogenized, and the supernatant stored at -20 °C until quantified. Briefly, microtiter polystyrene plates were coated with GAM IgM (1 μg/mL) and blocked with 1% BSA in PBS. Next, the IgM GXM binding mAb 2D10 (2 μg/mL) was added as a capture antibody [63], and the plate was incubated for 1 h. Culture supernatants were serially diluted on the plate and incubated for 1 h. The ELISA was completed by adding, in successive steps, mAb 18B7 (2 μg/mL) in buffer (PBS with 1% BSA), 1 μg of alkaline phosphatase-labeled GAM IgG1/mL in buffer, and 50 μl of *p*-nitrophenyl phosphate (5 mg/mL) in substrate buffer (0.001 M MgCl_2_ and 0.05 M Na_2_CO_3_, in 1 L [pH 9.8]). Between each step, the wells were washed with 0.05% Tween 20 in Tris-buffered saline. All incubations were done at 37°C for 1 h or 4°C overnight.

### WB analysis

WB analysis was conducted using cytoplasmic extracts from mouse brain cells made with a NE-PER nuclear and cytoplasmic extraction kit (TF). The mixture was centrifuged at 10,000 × g for 10 min at 4 °C, and the resulting protein content of the supernatant was determined using the Bradford method, employing a Pierce BCA protein assay kit (TF). Lysates were preserved in a protease inhibitor cocktail (TF) and stored at − 20 °C until use. Extracts were diluted with 2 × Laemmli sample buffer (Bio-Rad) and β-mercaptoethanol (Sigma). The mixture was heated to 90 °C for 5 min. Twenty-three µg of protein were applied to each lane of a gradient gel (7.5%; Bio-Rad). Proteins were separated by electrophoresis at a constant 130 V/gel for 90 min and transferred to a nitrocellulose membrane on the Trans-Blot Turbo Transfer System (Bio-Rad) at 25 V for 7 min. The membranes were blocked with 5% BSA in tris-buffered saline (TBST; 0.1% Tween 20) for 2 h at RT. A primary monoclonal claudin-5-, ZO-1-, occludin-, JAM-A-, RhoA-GTP, or RhoA (pan)-specific Ab (dilutions, 1:1000; Santa Cruz Biotechnology or Cell Signaling Technologies) was incubated overnight at 4 °C with TBST (5% BSA). After washing the membranes 3X with TBST for 10 min, a GAM IgG (H + L) conjugated to horseradish peroxidase was used as a secondary Ab (1:1000; Southern Biotech) and incubated with TBST (5% BSA) for 1 h at RT. The membranes were washed as described above. Protein bands were measured using the UVP ChemStudio imaging system (Analytik Jena) after staining each membrane with chemiluminescence detection reagents (TF). Quantitative measurements of individual band intensities in WB analyses for claudin-5, ZO-1, occludin, or JAM-A were performed using the NIH ImageJ software. Actin (dilution, 1:1,000; BD Biosciences), a cytoskeleton housekeeping protein, was used as loading controls to determine the relative intensity ratio. This WB protocol was previously described in [64] and modified accordingly for this study.

### Immunofluorescence confocal microscopy

Human Brain Endothelial Cells-5i (HBECs, American Type Culture Collection [ATCC]) were seeded at 0.75 × 10^5^ cells/glass coverslips in 24-well tissue culture plates and cultured for 24 h at 37°C and 5% CO_2_. Then, monolayers of cells were incubated without or with 10 μg/mL of GXM for 4 h at 37°C and 5% CO_2_. EDTA (10 μg/mL; Sigma) was used as a positive control. After each incubation, the solution with either GXM or EDTA was removed, and each coverslip fixed in a 4% paraformaldehyde solution for 10 min. Then, HBECs were rinsed 3X with PBS and blocked with PBS supplemented with 1% BSA followed by the addition of claudin-5-conjugated FITC and occludin-conjugated rhodamine binding mAbs (dilutions, 1:1000) in 1% BSA. The plate was then incubated for 1 h at 37°C. DAPI was diluted in PBS and used to stain HBEC nuclei for 1 h at RT. Slides were examined by confocal microscopy as described above and analyzed using NIH ImageJ software. Individual cells in each IF image were traced, the intensity of each protein determined, and the mean intensity per condition reported.

### BBB model

HBECs were seeded at 0.75 × 10^5^ cells/well in 0.4 μM pore transwells (Greiner Bio-ONE) and grown at 37 °C. Concurrently, human primary pericytes (Sciencell) were seeded at 1.5 × 10^5^ cells/well in 24-well tissue culture plates. After 24 h, transwells containing HBECs were transferred to wells containing pericytes and grown for 48 h.

#### a. Trans-Endothelial Electrical Resistance

Relative TEER of HBECs incubated with GXM (10, 50, or 100 μg/mL) for 5 h at 37 °C and 5% CO_2_ was recorded. HBECs incubated alone or with EDTA (10 μg/mL) were used as negative and positive controls, respectively. TEER was performed for 5 h using a CellZScope-E (nanoAnalytics). Fig. 8A illustrates the HBECs and pericytes BBB model used for the TEER measurements.

#### b. Streptavidin-HRP HBEC Barrier Permeability

Cells were stimulated with GXM for 1 h. Permeability of barriers was assayed by adding streptavidin-HRP (BD Biosciences) to the upper chamber and collecting samples from the bottom chamber after 30 min incubation at 37 °C and 5% CO_2_. Samples were treated with 3,3’,5,5’ tetramethylbenzidine (Sigma) and quenched with 2 M H_2_PO_4_ in a 96-well plate. Absorbance was measured at 450 nm, as described previously [65]. Fig. 8B depicts the BBB model used for the endothelial cell barrier permeability assays.

### Fungal transmigration studies

The *in vitro* BBB model consists of primary HBECs seeded at a density of 10^5^ in 5 μm pore transwells and grown for 24 h at 37 °C, 5% CO_2_. Then, we analyzed the passage of *C. neoformans* (10^5^ cryptococci) through the BBB *in vitro* for 4 h to determine the impact of GXM (10 µg/mL) on the integrity of the barrier. Untreated or EDTA (10 μg/mL)–treated tissue constructs were used as controls. CFU determinations after fungal transmigration were performed. Fig. 9A shows the HBECs BBB model used for the cryptococcal transmigration determination.

### Vascular reactivity to GXM

Isolated superior mesenteric arteries (SMA) were incubated in oxygenated Krebs buffer (130 mM NaCl, 14.9 mM NaHCO_3_, 4.7 mM KCl, 1.18 mM KH_2_PO_4_, 1.17 mM MgSO_4_-7H_2_O, 1.56 mM CaCl_2_-2H_2_O, 0.026 mM EDTA, 5.5 mM glucose, pH 7.4), with the perivascular fat carefully removed. SMA were cut into rings (2 mm in length) and cultured in vascular medium (ATCC) in an incubator at 37°C supplied by 5% CO_2_. SMA were treated *ex vivo* with 25 μg/mL of GXM for 40 min. The concentration of GXM was chosen based on results from TEER and HBEC barrier permeability experiments. A stock solution of GXM was made by dissolving the polysaccharide in distilled water and further diluted in Krebs solution. A vehicle consisting of distilled water was used as a control (untreated). After treatment, SMA rings were mounted in a Multi-Wire Myograph System 620M (Danish Myo Technology) for isometric tension recordings using a PowerLab 8/35 data acquisition system (ADInstruments Pty Ltd.). SMA rings were equilibrated in Krebs buffer for 30 min, and in chambers perfused with 5% CO_2_ in 95% O_2_ at 37°C, as described elsewhere [66]. In all experiments, SMA ring integrity was assessed by stimulation with 120 mM KCl (74.7 mM NaCl, 14.9 mM NaHCO_3_, 60 mM KCl, 1.18 mM, KH_2_PO_4_, 1.17 mM MgSO_4_-7H_2_O, 1.6 mM CaCl_2_-2H_2_O, 0.026 mM EDTA, 5.5 mM glucose). To test for the presence of endothelium, segments were contracted with 1 *µ*M phelynephrine (PE) (Sigma); once the vessels reached a stable maximum tension, the vessels were stimulated with 10 *µ*M acetylcholine (ACh) (Sigma), and relaxation was confirmed. For the stock solution, PE and ACh were dissolved in distilled water. SMA that achieved relaxation to ACh were considered to have a preserved endothelium. Cumulative concentration–response curves to ACh (1 nM to 10 *µ*M) and PE (1 nM to 10 *µ*M) were performed on intact SMA rings in the absence or presence of 25 μg/mL of GXM. Endothelium-dependent relaxation was recorded for ACh after maximal precontraction with 1 *µ*M PE.

### Statistical analysis

All data were subjected to statistical analysis using Prism 9.4 (GraphPad). Differences in survival rates were analyzed by the log-rank test (Mantel-Cox). *P* values for multiple comparisons were calculated by analysis of variance (ANOVA) and were adjusted by use of the Tukey’s post-hoc analysis. *P* values for individual comparisons were calculated using student’s or multiple *t*-test analyses. Concentration-response curve for vascular reactivity was log-transformed, normalized to percent maximal response, and fitted using a nonlinear regression. *P* values of <0.05 were considered significant.

## ACKNOWLEDGEMENTS

M.E.M., M.R.D., and L.R.M. were supported by the National Institute of Allergy and Infectious Diseases (NIAID award # R01AI145559) of the US National Institutes of Health (NIH). E. A. E. was supported by the National Institute of Mental Health (NIMH award # MH096625 and MH128082) of the US NIH, the National Institute of Neurological Disorders and Stroke (NINDS award # NS105584) of the US NIH, and The University of Texas Medical Branch internal funding. The funders had no role in the study design, data collection and analysis, decision to publish, or preparation of the manuscript.

## AUTHORSHIP CONTRIBUTIONS

All authors contributed to the project design and experimental procedures, analyzed data, provided the figure presentation, and manuscript writing.

